# Structural Models for a Series of Allosteric Inhibitors of IGF1R Kinase

**DOI:** 10.1101/2024.04.04.588115

**Authors:** Jyoti Verma, Harish Vashisth

## Abstract

The allosteric inhibition of Insulin-like Growth Factor Receptor 1 Kinase (IGF1RK) is a potential strategy to overcome selectivity barriers in targeting receptor tyrosine kinases. We constructed structural models of a series of 12 indole-butyl-amine derivatives which have been reported as allosteric inhibitors of IGF1RK. We further studied dynamics and interactions of each inhibitor in the allosteric pocket via all-atom explicit-solvent molecular dynamics (MD) simulations. We discovered that a bulky carbonyl substitution at the R1 indole ring is structurally unfavorable for inhibitor binding in the IGF1RK allosteric pocket. Moreover, we found that the most potent derivative (termed C11) acquires a distinct conformation, forming an allosteric pocket channel with better shape complementarity and interactions with the receptor. In addition to a hydrogen bonding interaction with V1063, the cyano derivative C11 forms a stable hydrogen bond with M1156, which is responsible for its unique binding conformation in the allosteric pocket. Our findings show that the position of chemical substituents at the R1 indole ring with different pharmacophore features influences molecular interactions and binding conformations of the indole-butyl-amine derivatives, hence dramatically affecting their potencies. Our results provide a structural framework for the design of allosteric inhibitors with improved affinities and specificities against IGF1RK.

## 1. Introduction

The type 1 insulin-like growth factor receptor (IGF1R) is a receptor tyrosine kinase (RTK) widely expressed in most somatic cells and tissues of all vertebrates [1]. IGF1R plays a critical role in cell growth, development, and differentiation [2]. IGFIR is a homodimer consisting of two *α* and two *β* chains connected by disulfide bonds [3]. It is synthesized as a precursor peptide of 180 kDa which undergoes posttranslational modifications to yield the mature *α*_2_*β*_2_ receptor [4,5]. The extracellular domain is comprised of the *α* chain and part of the *β* chain. The remaining *β* chain makes up the single-pass transmembrane domain and a cytoplasmic tyrosine kinase domain. Further, six different protein subdomains are identified within the extracellular region, three type III fibronectin domains designated FnIII-1, FnIII-2, and FnIII-3; an N-terminal leucine-rich domain (L1); a cysteine-rich domain (CR); and a second leucine-rich domain (L2) [5]. Unlike many other RTKs, IGF1R is activated by the extracellular binding of secreted growth factor peptides IGF-1 and IGF-2, resulting in autophosphorylation of the intracellular kinase domain and the formation of docking sites for signaling molecules [6].

As a growth mediator, IGF1R is ubiquitously expressed and promotes normal tissue development [7]. However, it has also been implicated in mitogenesis and suppression of apoptosis in tumor cells [8–10]. Various cancer types are associated with elevated levels of IGF1R and circulating IGF-1/IGF-2 [11,12]. Different mechanisms can influence IGF1R activity to produce cancer phenotypes, including, (i) IGF-1R overexpression caused by IGF1R gene aberrations, (ii) regulation of IGF-1R expression by modifier genes, (iii) increased levels of ligands, and (iv) transactivation by other membrane receptors [13]. Hence, various studies have demonstrated that IGF1R inhibition and downregulation cause substantial in vivo tumor cell death, impede tumorigenesis, and trigger a host response that eliminates surviving tumor cells [14–17]. Experimental investigations also suggest that inhibition of IGF1R kinase activity will implicate in disrupting tumor signaling leading to inhibition of tumor growth [17]. Therefore, IGF1R is an attractive target for chemotherapeutic interventions. Moreover, targeting RTK family of proteins is efficacious and well-tolerated, as demonstrated by the successful development of anti-tumor drugs against VEGFR, HER-2, and EGFR [18].

The therapeutics against IGF1R include antibodies to the extracellular domain that block receptor signaling and small molecule inhibitors of the tyrosine kinase domain. A recent analysis reported that, about 16 IGF1R drugs (both monoclonal antibodies and small molecule inhibitors) entered the oncology clinical trials between 2003 to 2021 [19]. However, none were approved for cancer treatment [19]. To date, several monoclonal antibodies against IGF1R such as ganitumab, figitumumab, dalotuzumab, dusigitumab, and xentuzumab have reached pre-clinical and clinical trials [20–24]. The small-molecule IGF1RK inhibitors have also shown considerable efficacy in pre-clinical trials [25]. The ATP competitive inhibitors, BMS-754807 and OSI906 (Linsitinib), are well studied although numerous trials indicated unsatisfactory outcomes [26,27]. AXL-1717 (picropodophyllo-toxin), a non-ATP competitive IGF1R kinase inhibitor has shown some potential in treating patients with relapsed malignant astrocytomas; nevertheless, its precise mode of action is yet unknown [28]. Many small molecule chemotypes still lack structural information and the mechanism of inhibition.

Despite persistent efforts to develop therapeutically useful IGF1R blockers of diverse molecular nature and mechanisms of action, the results of clinical trials have not been encouraging thus far. The most often cited explanations for the failure of IGF-1R-targeted treatments are crosstalk with other receptors, IR-IGF1R hybrid receptors, high degree of homology among the IGF1R and IR catalytic domains, and IR signaling compensation for IGF1R inhibition [25,29–32]. Hence, the specificity and selectivity of inhibitors continue to be further explored. Furthermore, allosteric inhibition has been considered as a possible strategy for increasing specificity and selectivity for kinase inhibitors [33,34]. However, experimental evidence of the allosteric interaction has been demonstrated only for a small number of allosteric kinase inhibitors [35]. Henrich et al. have described the design of a series of indole-butyl-amine derivatives as allosteric IGF1RK inhibitors and measured their biochemical activity against IGF1RK and IRK [36]. The allosteric binding site in the IGF1RK was characterized by X-ray crystallographic studies with one of the inhibitors (C10) from the series. However, the binding mode of other inhibitors in the series and their interactions with the IGF1RK remain unresolved. The inhibitor activity is invariably the consequence of various forces including hydrophobic interactions, polar interactions, shape and charge complementarity of the binding pocket and hence, it is challenging to interpret the differences in potencies of closely related compounds without a thorough structural analysis. Here, we report the structural model of each indole butyl amine derivative in complex with IGF1RK and detailed structural analysis to provide a rationale for the observed differences in the activities of these allosteric inhibitors. We performed a structural and energetic comparison of the interactions of each allosteric inhibitor using molecular docking, all-atom MD simulations, and free energy calculations. In addition, we report the unique binding conformation of the most potent derivative C11, and unravel the chemical basis for the design of potent allosteric inhibitors of IGF1RK.

## 2. Materials and Methods

### 2.1. Generating 3D Chemical Structures and Pharmacophore Features of Allosteric Inhibitors

We first obtained the chemical structures of the series of indole-butyl amine derivatives (labeled 1-12) reported as a novel class of allosteric IGF1RK inhibitors [36] and generated their atomic coordinates using the ChemAxon MarvinSketch package version 22.22 (https://www.chemaxon.com). The molecular structures were further categorized into four groups (Groups A-D) based on the ‘R’ group substituent (Figure 1). Furthermore, we generated the pharmacophore features for all inhibitors (1-12) using the Phase module of the Schrödinger software package [37,38]. The features included an H-bond donor (D), an H-bond acceptor (A), an aromatic ring (R), a hydrophobic site (H), a positive ionic (P) group, and a negative ionic (N) group.

**Figure 1.**
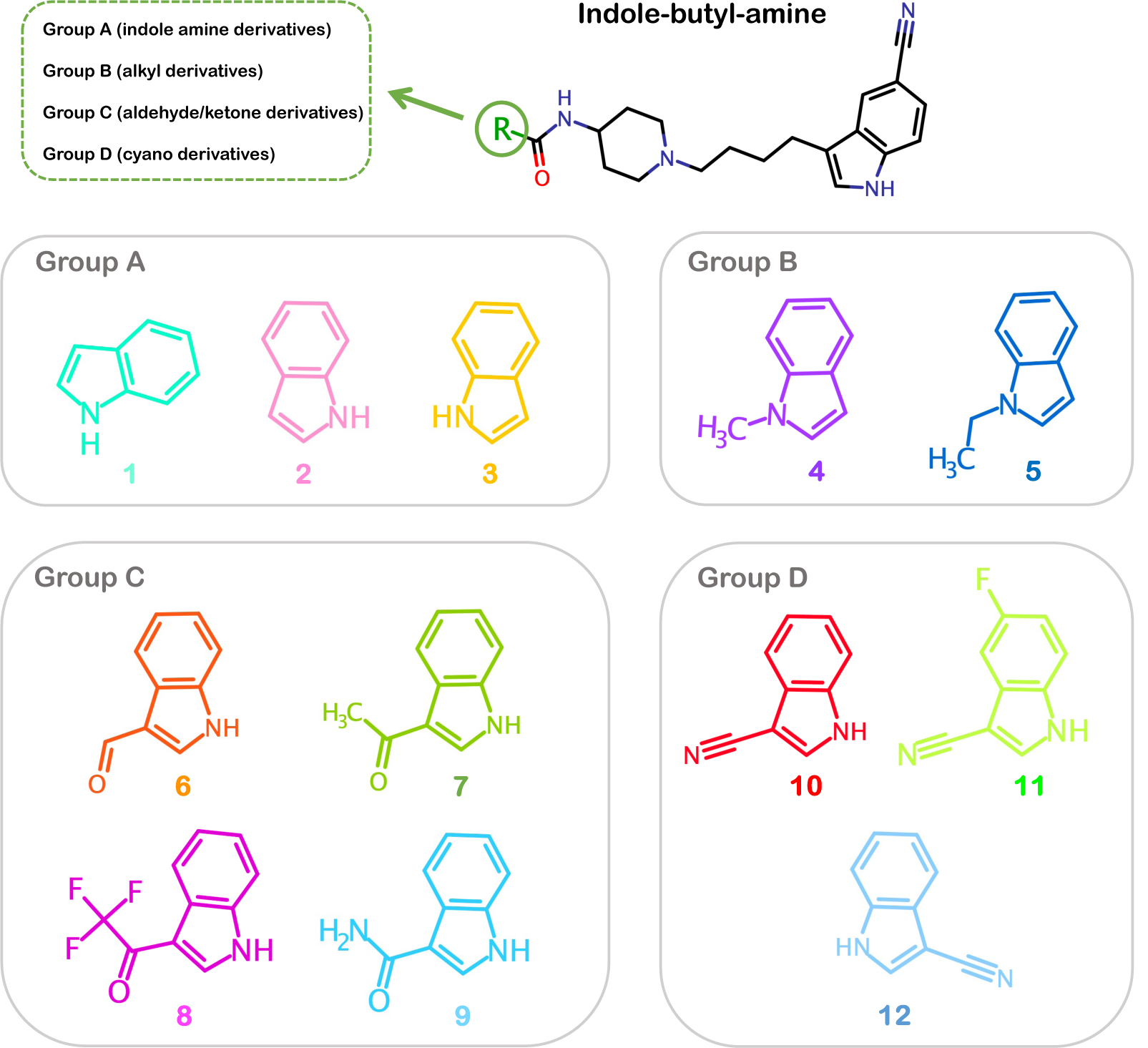
Chemical structures of the indole butyl amine derivatives. The derivatives are categorized into four groups (Groups A-D) based on their R-group substitutions. Group A: indole amine derivatives; Group B: alkyl derivatives; Group C: aldehyde/ketone derivatives; Group D: cyano derivatives.

### 2.2. Molecular Docking of Allosteric Inhibitors with IGF1RK

The crystal structure of IGF1RK in complex with the allosteric inhibitor C10 (PDB ID 3LW0) was obtained from the Protein Data Bank, which we used as a basis for our modeling of other inhibitor-docked structures [36]. We modeled the missing residues in the IGF1RK structure, A1097 through A1105 and G1169 through G1171, using the Modeller’s Chimera interface [39,40]. Eventually, we utilized the IGF1RK structure in complex with the inhibitor C10 for molecular docking of other inhibitors in the chemical series [36]. We optimized the structures of all inhibitors with correct molecular mechanics parameters before docking. The chemical structures were optimized using the OPLS3e force field with desalting and tautomer generation. The Epik module was used to generate all possible ionization states of the inhibitors at pH 7 *±* 2, and at most 32 stereoisomers were generated for each inhibitor. All these ligand optimization steps were accomplished using the LigPrep module of the Schrödinger software package [41,42]. Next, the protein structure was optimized using the protein preparation wizard of Schrödinger [41]. The protein was first pre-processed by adding missing hydrogen atoms, removing water molecules, and adding any missing sidechains in residues. Next, the H-bond optimization was performed to address any overlapping hydrogens added during the pre-processing. Finally, a restrained minimization was carried out with the OPLS3e force field to remove steric clashes while allowing the displacements of the heavy atoms up to 0.3 Å. Following minimization, the receptor grid generation panel was used to specify the protein as the receptor structure for docking and grid generation. A 20 Å^3^ receptor grid was generated around the IGF1RK allosteric pocket by selecting the bound inhibitor (C10) in the structure as the centroid of the grid. Finally, the docking of each inhibitor was performed in the extra precision (XP) mode of the Glide module [43,44]. The binding affinity (docking score) and the energetically favorable binding conformations of each inhibitor were determined [45]. From the XP docking, we obtained two poses for each inhibitor in the binding pocket. Ultimately, for each inhibitor, the docked conformation with the lowest docking score (more negative) was selected for further MD simulations. The docked conformations were the initial models of interactions of inhibitors in the binding pocket and further MD simulations were carried out to account for the flexibility of each inhibitor and the IGF1RK.

### 2.3. Molecular Dynamics (MD) Simulation Setup

We performed all-atom MD simulations of each IGF1RK-inhibitor complex using the GROMACS-2020.4 package [46]. The IGF1RK structure was parameterized using the AMBER99SB force field, while the inhibitors were parametrized with the general amber force field (GAFF) using the Antechamber module [47–49]. The protein and inhibitor coordinates were merged and the inhibitor topology information was integrated into the system topology file. Subsequently, a cubic simulation domain was generated with the protein at the center and at least 10 Å from the edge of the simulation domain. The TIP3P model for water molecules was used to solvate each system, and then Na^+^ and Cl*^−^* ions were added to neutralize each system’s net charge [50]. Following that, each system was energy minimized for 50000 steps using the steepest descent algorithm [51]. The inhibitor molecules were subjected to a positional restraint prior to unrestrained system equilibration. Each system was equilibrated first in the NVT ensemble for 10 ns at 300K using the velocity rescale thermostat followed by equilibration in the NPT ensemble for 50 ns using the Berendsen barostat at 1 atm pressure and 300 K temperature [52,53]. Finally, restraints were removed and a 200 ns production MD simulation was carried out for each IGF1RK/inhibitor complex. In all MD simulations, an integration time step of 2 fs was employed, and coordinates were saved every 100 ps.

### 2.4. Binding Free Energy Calculations

We further computed the binding free energy of each inhibitor for the IGF1RK domain to provide a distinct comparison of their binding modes. We estimated the enthalpy change and solvation free energy for each complex, using the Molecular Mechanics Generalized-Born Surface Area (MMGBSA) approach [54,55]. We used the gmxMMPBSA package which integrates the AMBER tools MMPBSA.py script for performing free energy calculations by utilizing MD trajectories [56,57]. We calculated the potential energy (bonded and non-bonded interactions), polar solvation energy, and non-polar solvation energy terms estimated over an average of 2000 frames in a 200 ns trajectory. The polar component was calculated utilizing the GB model with external and internal dielectric constants of 78.5 and 1.0, respectively. The non-polar solvation energy was estimated using the solvent-accessible surface area (SASA) approach, with the surface tension of the solvent set to 0.0072 and the surface tension offset set to 0.0. Furthermore, with per residue decomposition, the average contributions of residues to the binding free energy were also estimated.

### 2.5. Conformational and Interaction Analyses

The analyses of all MD trajectories were performed using the Gromacs built-in utilities from the Gromacs package (2020.4) and the Visual Molecular Dynamics (VMD) program [58,59]. We calculated the root mean squared deviation (RMSD) of the protein backbone atoms taking the initial structure as a reference and the root mean squared fluctuation (RMSF) was evaluated for the C*_α_* atom of each residue from its mean position. The RMSD was calculated for the heavy atoms of the inhibitor, taking the initial docked conformation as a reference. The kernel density distribution function in MATLAB was used to evaluate the RMSD distribution curve [60].

We performed the principal component analysis (PCA) to identify the essential conformational modes in the IGF1RK inhibitor complexes specifically at the allosteric pocket [61]. After building and diagonalizing a covariance matrix of the eigenvectors that represented the atomic variations in the protein and ligand atoms, a set of eigenvectors and their matching eigenvalues were obtained. The eigenvectors corresponding to essential movements in the protein-inhibitor complexes that have the greatest eigenvalues are called principal components, or PCs. We also constructed a free energy surface (FES) projected along the top two eigenvectors corresponding to the first and second principal components, PC1 and PC2, respectively.

All hydrogen bonds (H-bonds) were computed with a donor-acceptor interatomic distance threshold of 3.5 Å in VMD [59]. The protein-inhibitor interactions were determined using the LigPlot^+^ program [62]. All the images were rendered using the PyMOL software [63]. The MATLAB program was used to generate the plots [60].

## 3. Results and Discussion

### 3.1. Pharmacophore Modeling and Molecular Docking

In the current work, we elucidate the binding conformations of a series of 12 allosteric IGF1RK inhibitors (Table 1) while determining the rationale for differences in their potencies [36]. The series of indole butyl amine derivatives (C1-C12) have a large common substructure (1-[4(5-cyano-1H-indol-3-yl)-butyl]-piperidin-4-yl)-amide) with modifications in the indole ring located at one of the termini (Figure 1 and Table 1). We categorized these allosteric inhibitors based on the chemical moieties substituted at the R group (Figure 1). Group A constitutes the indole amine derivatives, C1, C2, and C3 which differ in the amide bond attachment on the indole ring. The compounds, C4 and C5, are alkyl derivatives in Group B with methyl and ethyl substituents added to the indolyl nitrogen. Group C includes the compounds C6, C7, C8, and C9 with more bulky aldehyde and ketone substituents. The cyano derivatives, C10, C11, and C12 are in Group D and have a cyano group attached to the indole ring.

**Table 1.**
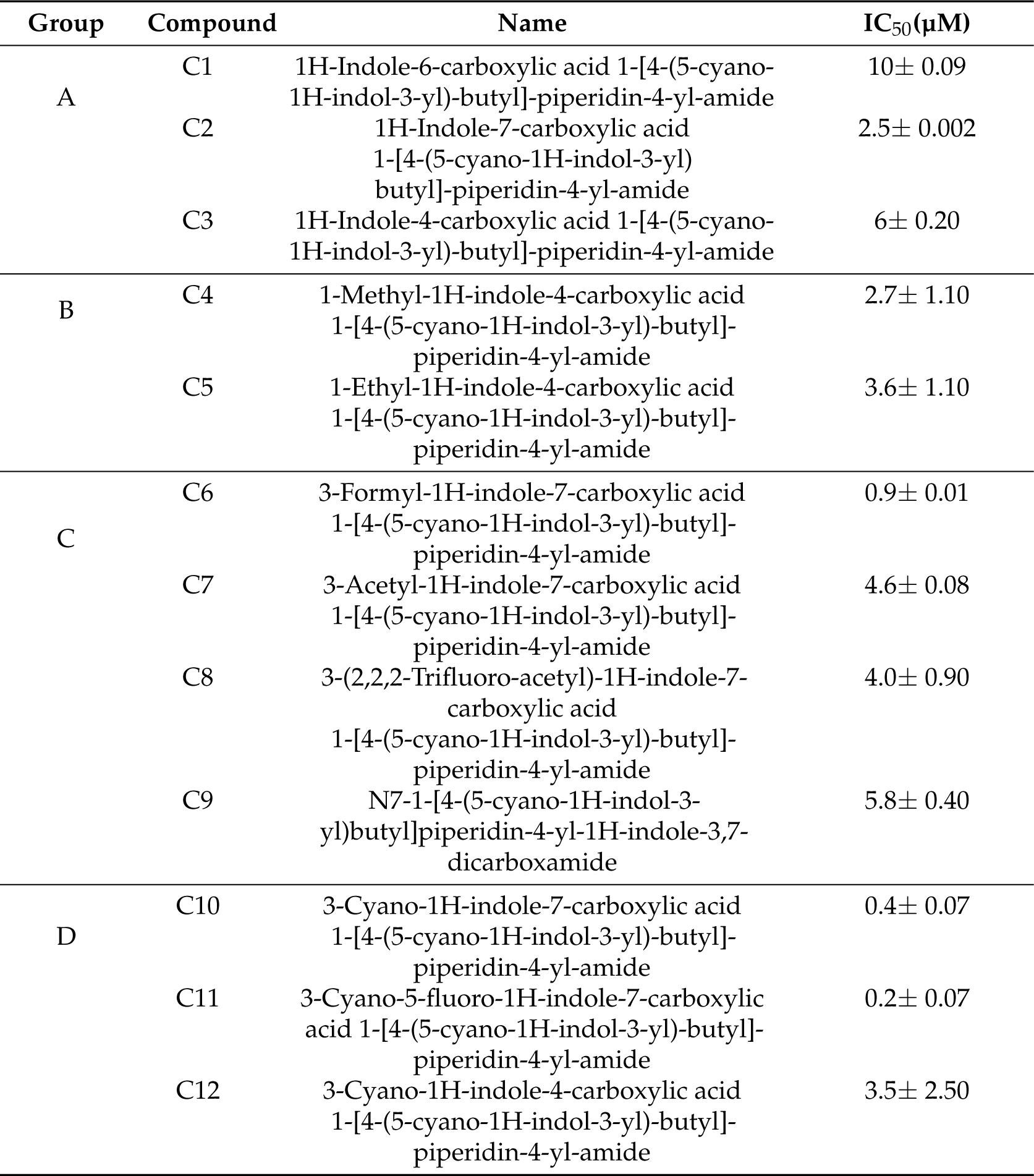
IGF1RK allosteric inhibitors with their respective IC_50_ (µM) values, as reported by Henrich et al. [36]. We categorized these derivates into four groups (Groups A-D) based on their ‘R’ group substituent.

| Group | Compound | Name | IC<sub>50</sub>(μM) |
| --- | --- | --- | --- |
| A | C1 | 1H-Indole-6-carboxylic acid 1-[4-(5-cyano-1H-indol-3-yl)-butyl]-piperidin-4-yl-amide | 10± 0.09 |
|  | C2 | 1H-Indole-7-carboxylic acid 1-[4-(5-cyano-1H-indol-3-yl)-butyl]-piperidin-4-yl-amide | 2.5± 0.002 |
|  | C3 | 1H-Indole-4-carboxylic acid 1-[4-(5-cyano-1H-indol-3-yl)-butyl]-piperidin-4-yl-amide | 6± 0.20 |
| B | C4 | 1-Methyl-1H-indole-4-carboxylic acid 1-[4-(5-cyano-1H-indol-3-yl)-butyl]-piperidin-4-yl-amide | 2.7± 1.10 |
|  | C5 | 1-Ethyl-1H-indole-4-carboxylic acid 1-[4-(5-cyano-1H-indol-3-yl)-butyl]-piperidin-4-yl-amide | 3.6± 1.10 |
| C | C6 | 3-Formyl-1H-indole-7-carboxylic acid 1-[4-(5-cyano-1H-indol-3-yl)-butyl]-piperidin-4-yl-amide | 0.9± 0.01 |
|  | C7 | 3-Acetyl-1H-indole-7-carboxylic acid 1-[4-(5-cyano-1H-indol-3-yl)-butyl]-piperidin-4-yl-amide | 4.6± 0.08 |
|  | C8 | 3-(2,2,2-Trifluoro-acetyl)-1H-indole-7-carboxylic acid 1-[4-(5-cyano-1H-indol-3-yl)-butyl]-piperidin-4-yl-amide | 4.0± 0.90 |
|  | C9 | N7-1-[4-(5-cyano-1H-indol-3-yl)butyl]piperidin-4-yl-1H-indole-3,7-dicarboxamide | 5.8± 0.40 |
| D | C10 | 3-Cyano-1H-indole-7-carboxylic acid 1-[4-(5-cyano-1H-indol-3-yl)-butyl]-piperidin-4-yl-amide | 0.4± 0.07 |
|  | C11 | 3-Cyano-5-fluoro-1H-indole-7-carboxylic acid 1-[4-(5-cyano-1H-indol-3-yl)-butyl]-piperidin-4-yl-amide | 0.2± 0.07 |
|  | C12 | 3-Cyano-1H-indole-4-carboxylic acid 1-[4-(5-cyano-1H-indol-3-yl)-butyl]-piperidin-4-yl-amide | 3.5± 2.50 |

Next, we identified the pharmacophore features of all compounds in the series, C1 to C12. This analysis revealed the following features: an H-bond donor (D), an H-bond acceptor (A), a hydrophobic site (H), an aromatic ring (R), and a positive ionic (P) group (Figure 2). Since the compounds have a large common substructure, the pharmacophore features defining the parent structure were identical among all inhibitors. The unique features of all compounds are highlighted in Figure 2. The presence of two aromatic rings was also consistent for all the R group substituents because of the common indole ring. The compounds from Group A (C1-C3), have an H-bond donor group besides two aromatic rings. The indole ring of the compounds C1, C2, and C3 has an amide group attached to it, which acts as an electron-donating substituent. On the other hand, the Group B compounds have a hydrophobic site with the presence of methyl and ethyl substituents on 4-indoyl nitrogen for C4 and C5, respectively. However, the compounds belonging to Group C have distinct pharmacophore features. For example, C6 and C7 have an H-bond donor as well as an acceptor group attached to their indole ring. The compound C8 has a more bulky substituent trifluoromethyl ketone, where the trifluoromethyl group is the hydrophobic site, the carbonyl oxygen is the H-bond acceptor, and the amide at the indole ring acts as the H-bond donor. Furthermore, C9 also has the carbonyl oxygen atom as the acceptor and the 7-indolyl as the donor, as well as two additional H-bond donor groups from the carboxylic amide substituent. The cyano derivatives (Group D) C10 and C12 are isomers having both H-bond donor and acceptor groups. Besides a donor and an acceptor group, C11 has an additional hydrophobic site with a fluorine atom attached to the indole ring at the 5th position. Following the chemical characterization, we further compared these allosteric inhibitors structurally and energetically in the IGF1RK allosteric pocket.

**Figure 2.**
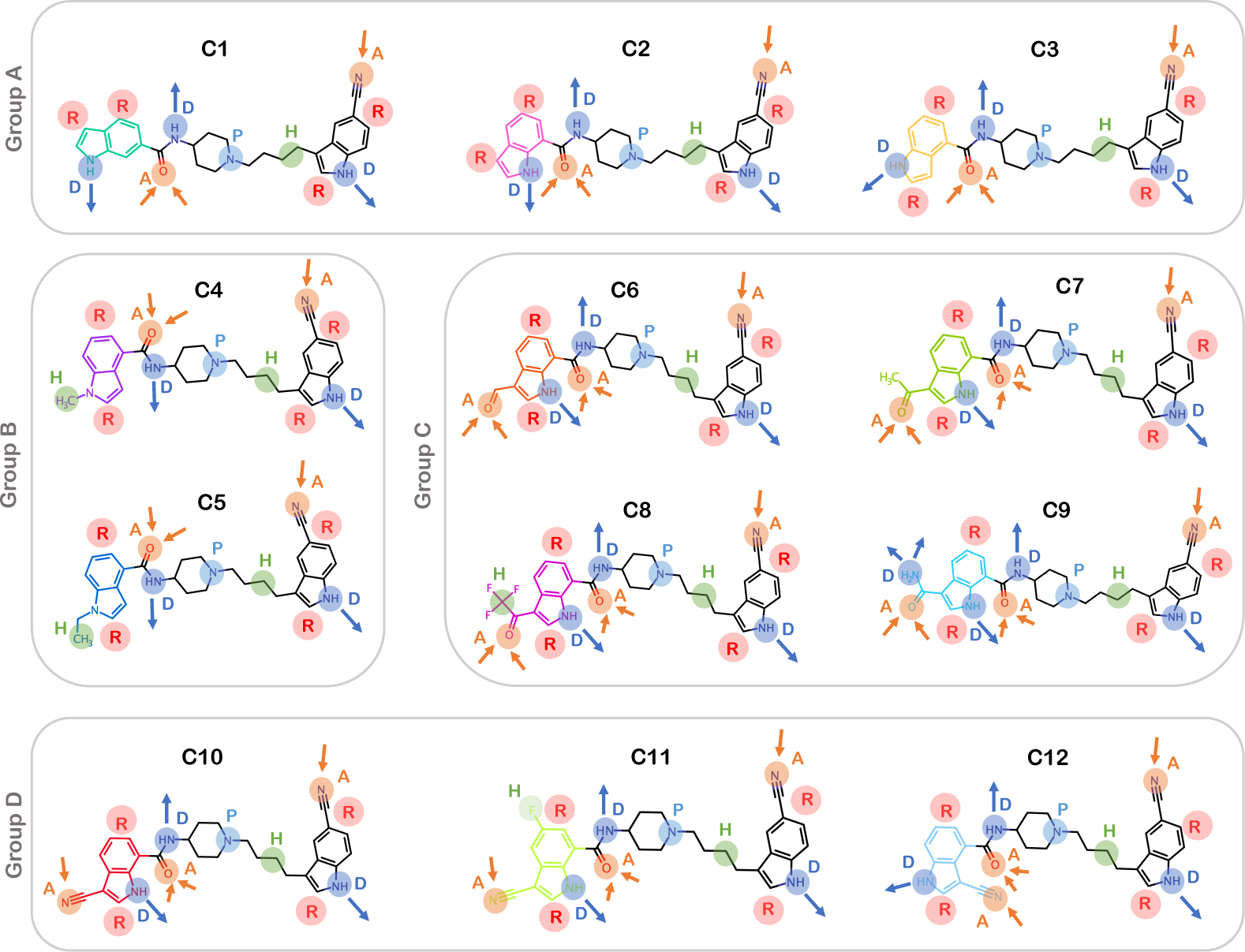
Pharmacophore features in each of the derivatives. Each feature is represented by a unique color and alphabet. A: Acceptor; D: Donor; H: Hydrophobic; P: Positive ionic; R: Ring.

Using the molecular docking approach, we initially modeled and predicted the conformation of each allosteric inhibitor in the IGF1RK allosteric site. We used the X-ray crystal structure of the IGF1RK domain in complex with the allosteric inhibitor C10 (PDB ID 3LW0) as the basis for the initial modeling of remaining 11 inhibitors. For each inhibitor, the conformation with the lowest docking score was selected. We further compared the docked conformations of all 11 derivatives with the crystallized conformation of C10 in the IGF1RK allosteric pocket (Figure 3). Overall, the conformation of the common substructure of these derivatives attained a similar configuration. All compounds showed an identical configuration of the 5-cyano indole ring (R2), while the conformational differences were observed along the R substituent at the other end (R1) (Figure 3B). The docking results reflect that the hydrogen bonding with the carbonyl oxygen atom of V1063 could be a significant factor for inhibitor binding to the IGF1RK allosteric pocket. Hence, an H-bond donor group at the 5-cyano indole ring (R2) is a key feature of an allosteric inhibitor candidate for IGF1RK. The electron-withdrawing cyano substituent at the R2 indole ring potentially also has a critical role in interaction with residues in the *α*C-helix. The presence of these chemical features at the R2 indole ring (5-cyano indole) resulted in its identical conformation in all inhibitors at the IGF1RK pocket. To corroborate these observations and further investigate the significance of chemical modifications at the R1 indole in the inhibitor potency, we studied the conformational dynamics of each complex using all-atom MD simulations.

**Figure 3.**
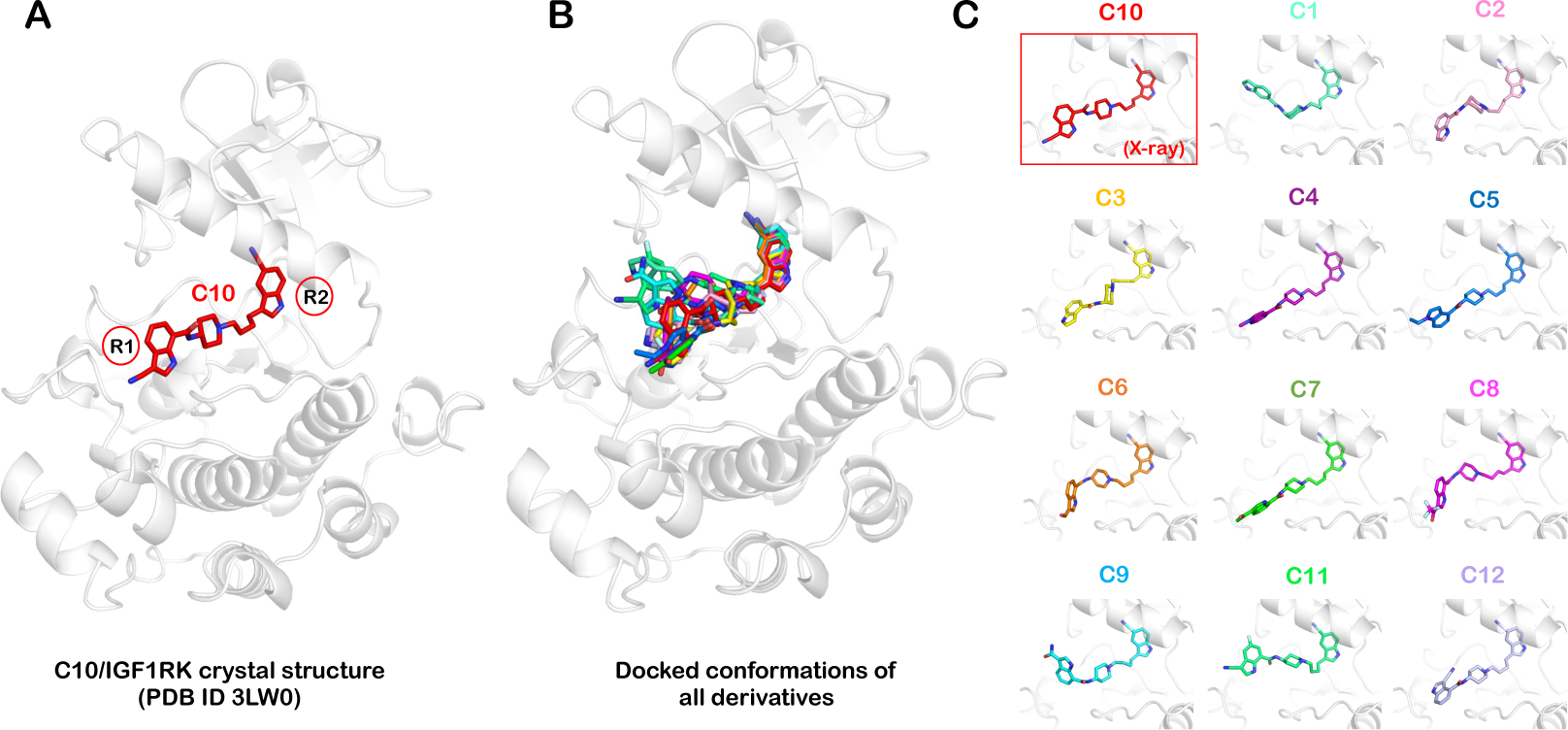
Docked conformations of allosteric inhibitors in the IGF1RK binding pocket. **(A)** The X-ray crystal structure of the C10 inhibitor bound to the IGF1RK. **(B)** Structural superimposition of the docked conformations of 11 indole butyl amine derivatives on the X-ray crystal structure of IGF1RK bound to the C10 derivative. **(C)** The docked conformation of each inhibitor in the IGF1RK pocket.

### 3.2. Conformational Dynamics of Complexes of IGF1RK/Inhibitors

We performed MD simulations for each complex to understand the conformational changes in the IGF1RK structure upon inhibitor binding in the allosteric pocket. In parallel, the dynamics of each allosteric inhibitor were studied to comprehend differences in their binding affinities. We first computed the RMSD of the IGF1RK backbone atoms and the non-hydrogen atoms of each inhibitor. For Group A inhibitors, C1, C2, and C3, the IGF1RK backbone RMSD distribution curves with single peaks between 2.0 Å and 3.5 Å (Figure S1A). The protein backbone RMSD for the C2 complex has a narrow distribution curve that shifts further to the left, indicating a lower RMSD value. The Group B inhibitor C4 has an average protein backbone RMSD value of 2.5 Å, whereas C5 has a broader distribution curve ranging between 2 Å and 4.5 Å, reflecting higher deviations in the protein backbone atoms (Figure S1A). The Group C inhibitors C6, C7, C8, and C9 have relatively higher IGF1RK backbone RMSD values, with wider distribution curves along the RMSD axis (Figure S1A). Further, in Group D, the cyano derivative C10 has the IGF1RK backbone RMSD distributed between 1.5 Å and 4.5 Å. In contrast, its isomer C12 has the lowest protein backbone RMSD of about 1.9 Å. Also, the inhibitor C11, on the other hand, has a IGF1RK backbone RMSD value of 2.2 Å. Overall, the IGF1RK backbone was more flexible with Group C inhibitors and was relatively less flexible with Group A and Group D inhibitors.

We further report the mean RMSD value derived for the non-hydrogen atoms of each inhibitor (Figure 4). Among Group A inhibitors, C2 has the lowest mean RMSD value of 0.8 Å, followed by C3 and C1 with 1.8 Å and 2.3 Å, respectively (Figure 4). Also, the RMSD distribution curve for C2 spans between 0.5 Å and 1.5 Å (Figure S1B). However, the inhibitors C1 and C3 have their RMSD values distributed between 1 Å and 3.5 Å, and 1 Å to 3 Å, respectively (Figure 4 and Figure S1B). These observations suggest that the inhibitor C2 (with the difference in the amide position at the indole ring R1 compared to C1) has lower deviations, leading to a relatively more stable conformation (Figure 4 and Figure S2). The mean RMSD for the C4 inhibitor is 0.86 Å, whereas for C5, it is 1.67 Å (Figure 4). Although C4 shows a single peak and a narrow distribution, C5 showed a distribution with two peaks, spanning RMSD values between 1 Å and 2.5 Å (Figure S1B). Hence, in Group B, the inhibitor C4 (methyl substituted) exhibits overall low deviations in the complex. For Group C inhibitors, C6, C7 and C8, the mean RMSD are 1.6 Å, 0.8 Å, and 1.6 Å, respectively.

**Figure 4.**
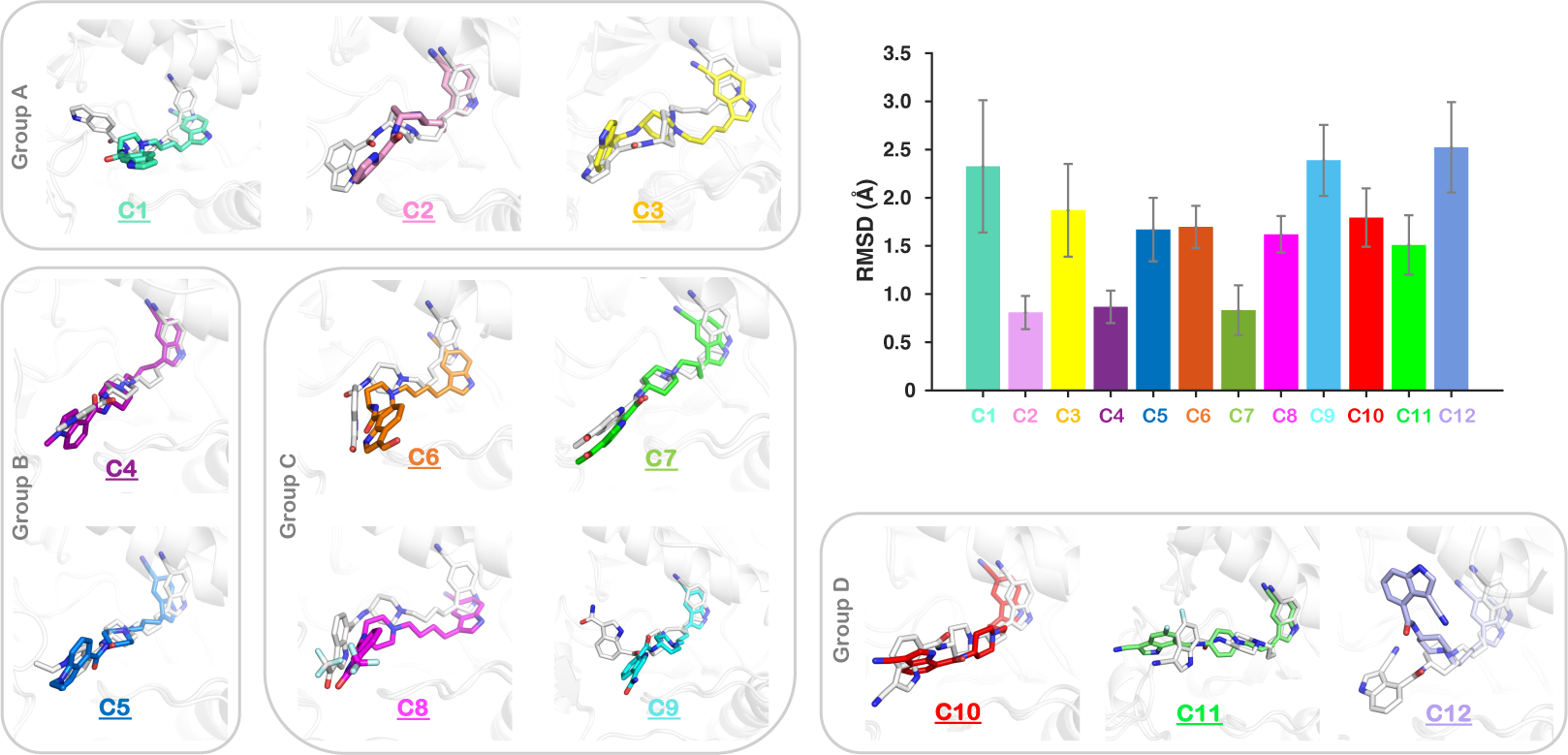
RMSD data for the heavy atoms of each inhibitor. The snapshot for each inhibitor with their initial docked conformation (white) superimposed on the most populated conformation in MD simulations is shown.

The restricted movement/flexibility of the substituted ketone and aldehyde groups of these inhibitors in the allosteric pocket may account for their lower RMSD values (Figure 4). However, the inhibitor C9 has an RMSD value of 2.3 Å, reflecting relatively more flexibility of the inhibitor in the allosteric pocket. Among the cyano derivatives (Group D), C10 (previously co-crystalized with IGF1RK) has the mean RMSD value of 1.7 Å. In contrast, its isomer C12 has the highest inhibitor RMSD value of 2.5 Å. The inhibitor C12 is highly flexible in the allosteric pocket and deviates from the allosteric region during simulations (Figure 4 and Figure S2). The inhibitor C11, on the other hand, has an RMSD value of 1.5 Å, which is the lowest among Group D compounds. These findings altogether imply that the position of the amide, as well as the cyano group or any electron-withdrawing group on the indole ring, has a critical role in inhibitor stability.

We also evaluated the RMSF for the IGF1RK residues to characterize their dynamics. The overall trend of residue fluctuations in all complexes was relatively similar. There was an evident fluctuation peak corresponding to the loop region (residues 1093 to 1115) with the RMSF values ranging between 4 Å and 6 Å in all complexes (Figure S1C). The fluctuation amplitude corresponding to the RMSF of the catalytic loop (residues 1133 to 1142) and the activation loop (residues 1152-1174) residues are comparable for all inhibitor complexes. Residues from these loops also specifically interact with the substituted R group of the inhibitors. The time evolution trace of the hydrogen bonds formed between the allosteric inhibitors and IGF1RK is shown in Figure S1D. The maximum number of hydrogen bonds that may form is 4-5; however, throughout the trajectory, all inhibitors consistently only display a small number (1-2) of hydrogen bonds. We further elucidated the stability of these hydrogen bonding interactions in each complex. We also measured the radius of gyration (R*_g_*) for each complex to indicate the compactness of the protein structure. We observed that the mean R*_g_* value for all complexes was in the range between 19.7 Å and 20.3 Å (Figure S3). The consistent value of R*_g_* for all complexes indicates that the IGF1RK structure remained compact upon binding to inhibitors. Although a visual inspection of simulation trajectories provided insight into the binding conformations of the allosteric inhibitors in the IGF1RK pocket (Figure S2), we further investigated the predominant binding conformation for each inhibitor.

### 3.3. Dominant Conformation of Each Inhibitor in the Allosteric Pocket

We performed the principal component analysis of all MD trajectories to understand the key conformational motions and extract the structural information for each IGF1RK-allosteric inhibitor complex. The first few principal components define the collective motions of the localized atomic fluctuations. The two-dimensional FES plots of each complex for the first two principal components (PC1 and PC2) are represented in Figure 5. The conformational phase space and the energy basins for each complex were unique and different. From the free energy maps, we identified a global minimum conformation of each inhibitor that is associated with the lowest free energy. The allosteric inhibitors C1 and C2 from Group A and C4 and C5 from Group B have two well-separated minima with higher energy barriers (Figure 5). Hence, conformations corresponding to both minima were analyzed for these inhibitors. Despite a widespread conformational phase space, a single global free energy minimum was observed for Group C and Group D inhibitors. The inhibitors C6 and C7 have well-separated global minima, while C8 and C9 have multiple local minima and a single global minimum with lower energy barriers. The cyano derivatives (Group D) on the other hand have a single global free energy minimum (Figure 5). We further obtained and evaluated the conformations of all allosteric inhibitors with IGF1RK corresponding to these minima.

**Figure 5.**
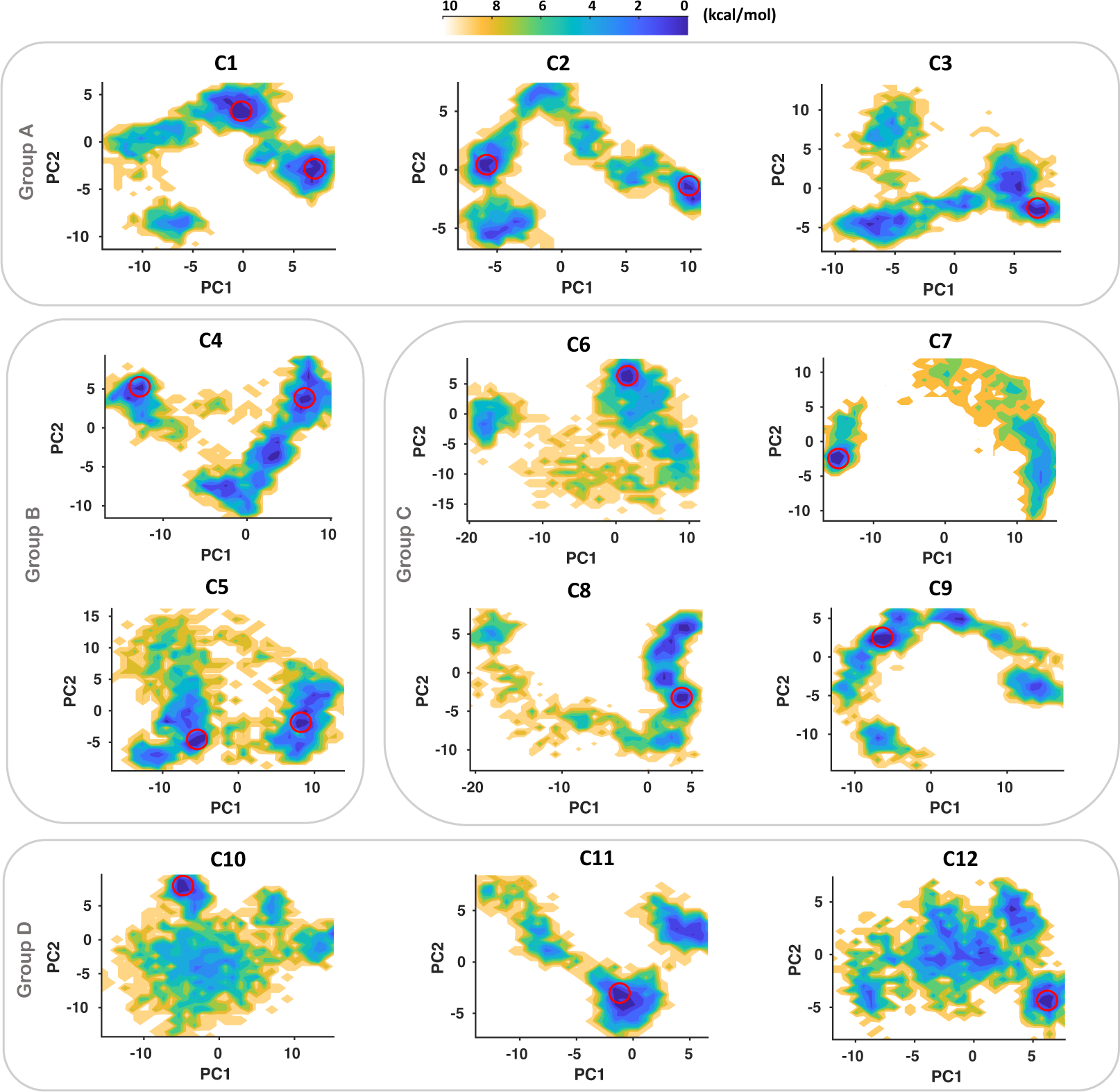
The free energy surface (FES) projections along two principal components (PC1 and PC2) for each inhibitor complex with IGF1RK. The minima corresponding to the lowest free energy in each plot are marked by red circle(s). The free energy is given in kcal/mol and indicated by the color palette from lower (blue) to higher (yellow) values.

We assessed the predominant conformation of each of the allosteric inhibitors in the IGF1RK allosteric pocket. The structural superimposition of all 12 complexes is shown in Figure 6. An overall assessment of the binding modes of all allosteric inhibitors showed that the Group A (C1-C3), B (C4-C5), and C (C6-C9) derivatives occupy similar configurations clustered at the same region of the allosteric pocket. As shown in Figure 6A, the shared indole R2 moiety in all inhibitors has similar conformations. Furthermore, the local conformational changes of the A-loop and the *α*C-helix caused by inhibitor binding, affect the IGF1RK allosteric pocket characteristics. As a consequence, Group D, cyano derivatives, C11 and C12, protrude away from the proximal allosteric binding pocket region and attain different conformations with respect to their initial docked conformations (Figure 4 and Figure 6). A detailed assessment of the conformations of these derivatives is discussed in the following section.

**Figure 6.**
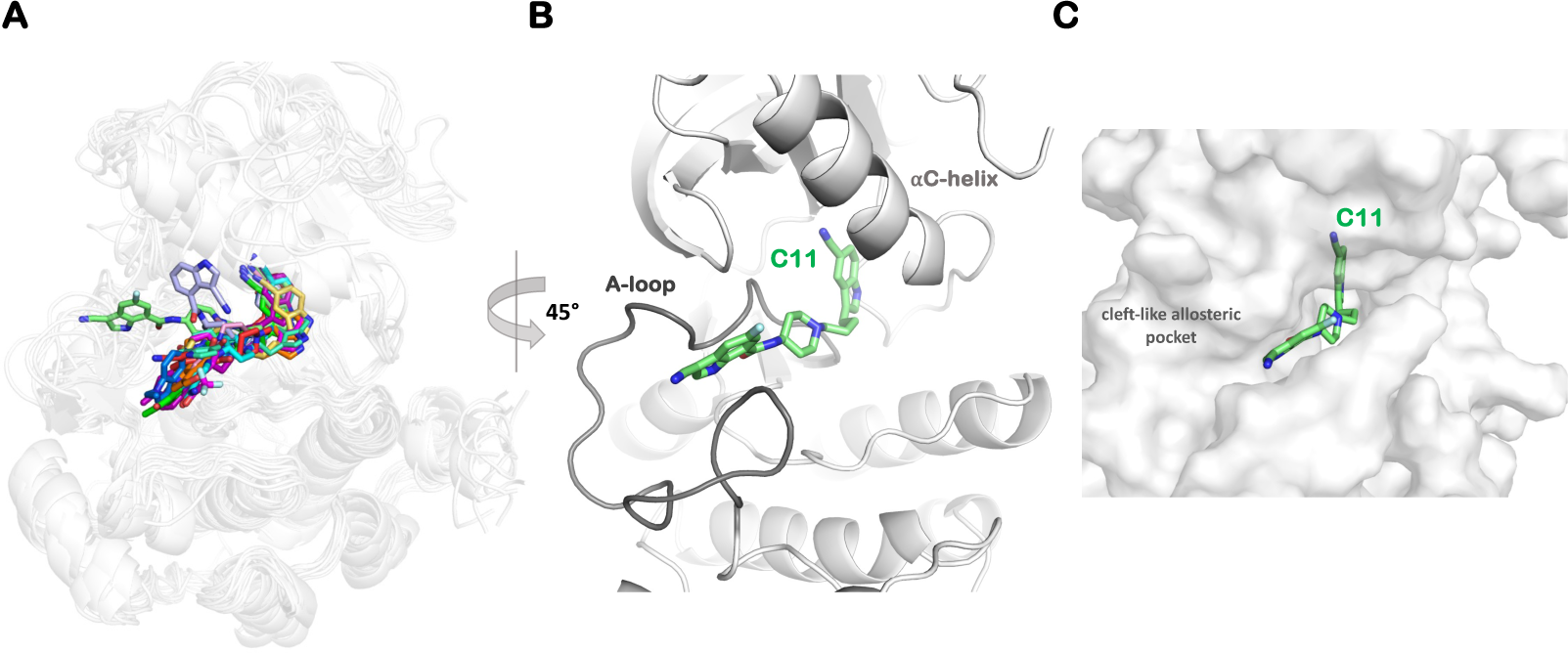
Binding conformation of the indole butyl amine derivatives (C1-C12) in the allosteric pocket of the IGF1RK. **(A)** Structure superposition of inhibitor conformations corresponding to the lowest free-energy minimum (see Figure 5) for each IGF1RK-inhibitor complex. **(B)** The unique binding mode of the most potent derivative C11 in the IGF1RK allosteric pocket. **(C)** A surface view of the binding pocket with the C11 derivative.

### 3.4. Effect of Substitutions at the R1 Indole Ring and Unique Binding Mode of C11

The effect of chemical substitutions in indole-butyl-amine derivatives on their biochemical activities (IC_50_) against IGF1RK was investigated by Henrich et al. [36]. We further recapitulate the substitutions at the R1 indole ring with molecular and structural aspects. The comparison of the whole series of derivatives emphasizes the molecular basis for the activity differences among them. We observed a favorable H-bond interaction between the acceptor carbonyl oxygen atom of V1063 and the donor group NH9 of the R2 indole ring in all inhibitor complexes (Figures S4, S5, S6 and S7). This interaction, however, was disrupted in the complexes of C1, C6, C8, and C9, but it was stable and consistent for all other inhibitors (Figure S4 and Figure S6). Furthermore, the atoms of the R2 indole ring participate in interaction with the hydrophobic residues V1063, M1054, and M1079, and these interactions are conserved in all complexes (Figure S9). This suggests that the 5-cyano indole (R2) at one end of the inhibitor establishes a stable hydrogen bond and is buried in a hydrophobic cavity created by M1054 and M1079, making it suitable for the IGF1RK allosteric pocket. Hence, we show that an electron-donating group with two aromatic rings is responsible for crucial interactions with the 5-cyano indole (R2) in the binding pocket. The substitution at the R1 indole ring, however, accounts for the differences in their affinities for IGF1RK. Residues H1133, R1134, I1151, G1152, D1153, and L1174 are common residues involved in interactions with the R1 indole ring of each derivative. In particular, the sidechain of the residue R1134 interacts with the R1 indole ring, influencing its conformation. This cation-*π* interaction with R1134 is mostly conserved in all inhibitors. For C1, the position of the amide at the indole ring is disfavoring its initial interactions at the binding pocket, resulting in dissociation of the H-bond with V1063 while forming new H-bonds with K1171 and G1172 (Figure S4). Furthermore, the dissociation of H-bond in C6, C8, and C9 could be due to the inability of the bulky carbonyl groups like ketone, aldehyde, and amide to fit into the allosteric pocket (Figure S6). Further, the cyano derivative C12 interacts with the *α*C-helix by extending out of the allosteric region. This observation emphasizes that the position of the cyano group in C12 might cause steric hindrance and, hence, is difficult to accommodate in the allosteric pocket (Figure S7). Altogether, the position of each substitution at the R1 moiety is critical as it is responsible for both electronic properties and steric effects in the structure. The R1 indole ring with an electron donating group (amide group) at position 7 and an electron acceptor group at position 3 likely favors the inhibitor binding as well as its activity against IGF1RK.

We performed a comprehensive analysis of the binding free energy using the MMGBSA approach to describe the molecular interactions and energetics of binding of allosteric inhibitors. The total binding free energy (ΔG) of each allosteric inhibitor with the IGF1RK is shown in Figure S8. The time evolution traces for the binding energy estimated at each frame, indicate a relatively stable curve for the cyano derivatives, C10 and C11 (Figure S8). The ΔG*_bind_* for individual residues involved in the interaction with each derivative is also given in Figure S9. The C10 and C11 compounds are the most potent derivatives reported against IGF1RK, with IC_50_ values of 0.4 *µ*M and 0.2 *µ*M, respectively (Table 1). The derivative C10 has been co-crystallized with the IGF1RK domain with the most favorable conformation of the 3-cyano indolyl carboxamide substituent (R1) in the X-ray structure. Our structure determined by analyzing ensemble conformations of C10 has its 3-cyano indolyl carboxamide substituent (R1) stacked parallel to the C-loop, forming favorable cation-*π* interactions with the residue R1134 (ΔG*_bind_* = *−*2.32 kcal/mol) (Figure 7). Further, the IGF1RK residues M1054, V1063 and M1079 significantly contribute to the binding of C10 with ΔG*_bind_* values of *−*2.45 kcal/mol, *−*1.67 kcal/mol and *−*1.56 kcal/mol, respectively (Figure 7).

**Figure 7.**
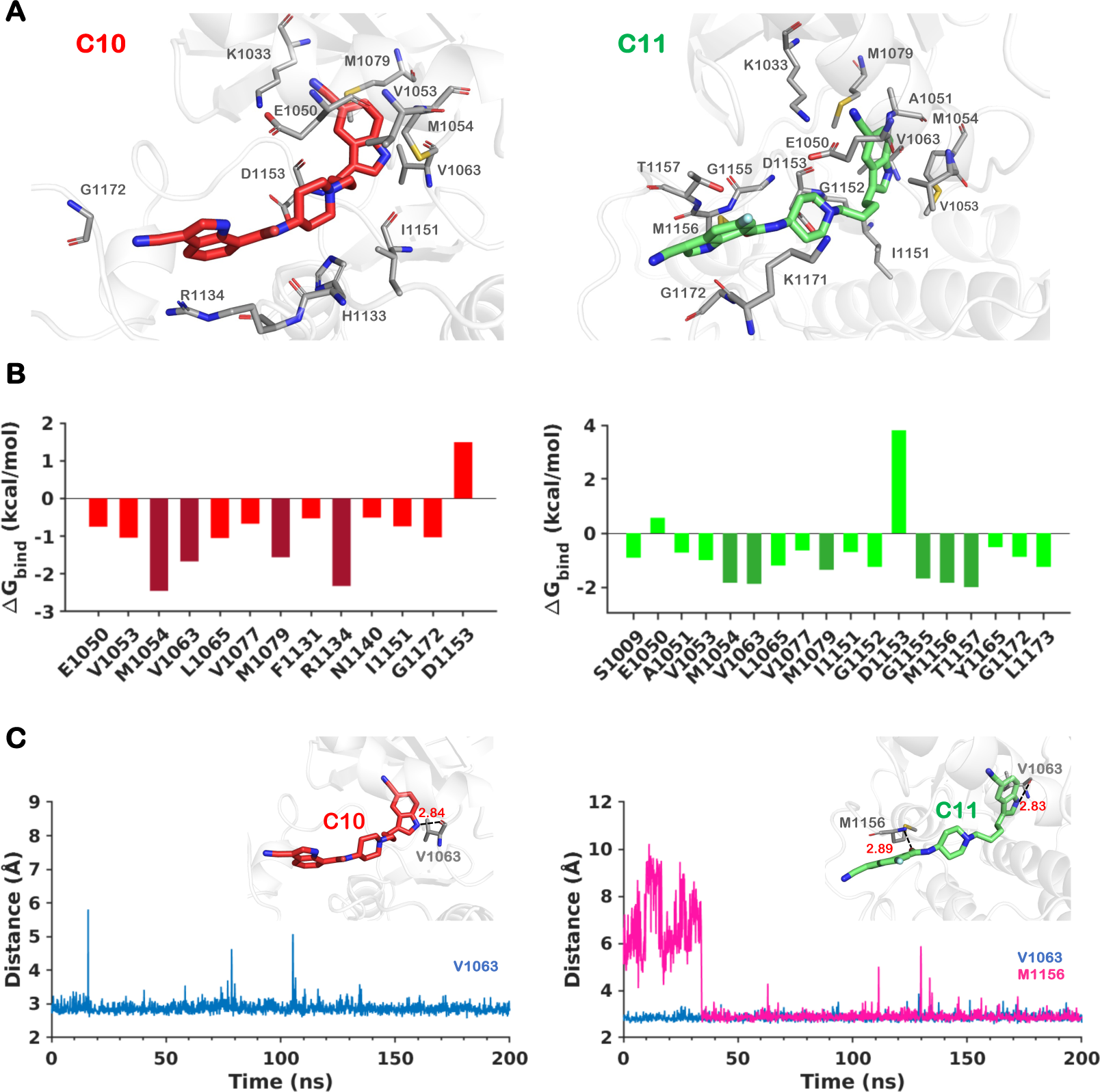
The binding conformations and the interactions of the allosteric inhibitor C10 and C11 in the IGF1RK pocket. **(A)** The binding mode of C10 and C11 in the allosteric pocket with the residues involved in the interactions. **(B)** The residue-level energy contribution (ΔG*_bind_*) for both complexes. The residues having higher/maximum energy contributions are represented in darker shades. **(C)** The distance traces for the unique hydrogen bonds between the IGF1RK residues and inhibitor atoms.

Our findings show that the most potent C11 derivative achieves a unique conformation in the IGF1RK pocket compared to all other derivatives. Figure 6B shows that C11 induces a conformational change in the A-loop that transforms the shape of the allosteric pocket in IGF1RK and offers a cleft-like formation at the allosteric binding region. A detailed inspection of these two compounds bound to IGF1RK shows that the binding of C11 is influenced by an additional hydrogen bond with the residue M1156, providing better shape complementarity and interactions with the receptor (Figure 6 and Figure 7). The indole ring R1 of the C11 is stacked between the sidechains of residues K1171 and T1157 and makes favorable interactions with residues G1155 and G1172 (Figure 7). Hence, the R1 substituent of C11 is stablized by the favorable energetic contributions of these residues, with G1155 (ΔG*_bind_* = *−*1.68 kcal/mol), M1156 (ΔG*_bind_* = *−*1.84 kcal/mol), and T1157 (ΔG*_bind_*= *−*2.00 kcal/mol) contributing most to the binding free energy of C11 in the IGF1RK pocket (Figure 7). The presence of a fluorine atom (hydrophobic site; H) at the indole ring provides improved activity by supporting and directing a stable conformation of the R1 indole ring in a cleft-like formation of the A-loop, while the carbonyl oxygen (acceptor; A) secures a stable hydrogen bond with the donor group at M1156. We thereby propose a structural rationale at the molecular level behind the higher potency of the cyano derivative C11.

## 4. Conclusions

We used pharmacophore modeling, molecular docking, and all-atom MD simulations to structurally rationalize the structure-activity relationships and binding affinities for a series of 12 allosteric inhibitors of IGF1RK. Our results provide structural explanations for the experimentally-observed activities of the indole-butyl-amine derivatives against IGF1RK. We show that the key interaction with the V1063 residue was initially preserved in all complexes, with a network of hydrophobic interactions involving the sulfur side chains of the residues M1054 and M1079. Hence, the presence of an electron-donating group with two aromatic rings of the 5-cyano indole of the inhibitor is essential for these interactions. Our findings suggest that bulky carbonyl substitutions such as ketone, amide, or aldehydes at the R1 indole ring are less favorable for accommodating inhibitors in the IGF1RK allosteric pocket. The position of the electron-withdrawing (acceptor; A), electron-donating (donor; D), and hydrophobic (H) substituents at the R1 indole ring influence the binding conformations of allosteric inhibitors, impacting the IGF1RK inhibition. Furthermore, we show that the binding of C11 induces the formation of a distinctive pocket with the A-loop and forms a unique H-bond with M1156. This allosteric pocket channel has a striking shape as well as chemical complementarity for the inhibitor C11. Thus, our results explain the reasoning underlying the higher potency of the cyano derivative C11. These structural investigations offer molecular explanations for variations in potencies of allosteric inhibitors, emphasizing the conformational changes and molecular interactions that can be exploited in the design of further improved allosteric inhibitors of IGF1RK and other kinases.

## Supplementary Materials

The following supporting information can be downloaded at: https://www.mdpi.com/article/10.3390/ijms1010000/s1

## Author Contributions

Conceptualization, J.V and H.V; methodology, J.V; software, J.V.; validation, J.V; formal analysis, J.V.; investigation, J.V; data curation, J.V; visualization, J.V; writing—original draft preparation, J.V; writing—review and editing, H.V; supervision, H.V; resources, H.V; project administration, H.V; funding acquisition, H.V. All authors have read and agreed to the published version of the manuscript.

## Funding

This research was funded by the National Institutes of Health (NIH) through Grant Nos. R35GM138217 and P20GM113131.

## Data Availability Statement

The data that support the findings of this study are available in the supporting information of this article.

## Acknowledgments

We acknowledge the financial support provided by the National Institutes of Health (NIH) through Grants R35GM138217 and P20GM113131. The content is solely the responsibility of the authors and does not necessarily represent the official views of the NIH. We are grateful for computational support through Premise, a central shared HPC cluster at UNH supported by the Research Computing Center.

## Conflicts of Interest

The authors declare no competing financial interest

## Abbreviations

The following abbreviations are used in this manuscript:

CR: Cysteine-rich domain
EGFR: Epidermal Growth Factor Receptor
HER-2: Human Epidermal Growth Factor Receptor 2
IGF1R: Insulin-like Growth Factor 1 Receptor
IGF-1: Insulin-like Growth Factor 1
IGF-2: Insulin-like Growth Factor 2
IR: Insulin Receptor
RTK: Receptor Tyrosine Kinase
VEGFR: Vascular Endothelial Growth Factor Receptor

## Disclaimer/Publisher’s Note

The statements, opinions and data contained in all publications are solely those of the individual author(s) and contributor(s) and not of MDPI and/or the editor(s). MDPI and/or the editor(s) disclaim responsibility for any injury to people or property resulting from any ideas, methods, instructions or products referred to in the content.

## Notes

### Competing Interest Statement

The authors have declared no competing interest.

### Summary of Updates

Figure S10 was cited in the main manuscript.

