## Supporting Information for "Structural Models for a Series of Allosteric Inhibitors of IGF1R Kinase"

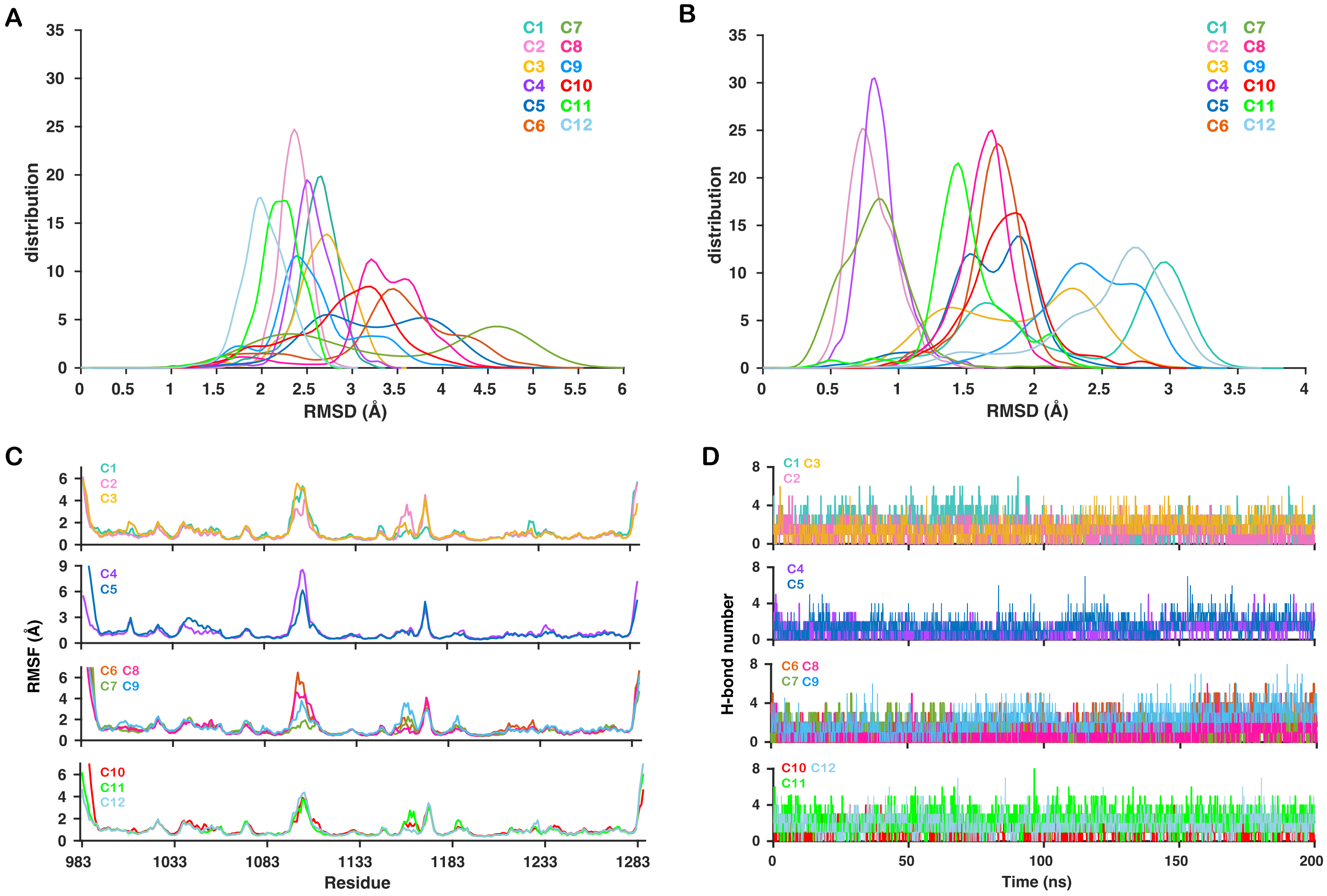


**Figure S1: The metrics for characterizing dynamics of each IGF1RK inhibitor complex.** (A) The Root Mean Squared Deviation (RMSD) distributions for the protein backbone atoms in each complex. (B) The RMSD distributions for the heavy atoms of each inhibitor. (C) Root Mean Squared Fluctuation (RMSF) per residue for the Cα atoms for each complex. (D) The time evolution traces of the number of hydrogen bonds between the inhibitor and the IGF1RK residues. A cutoff distance of 3.5 Å was used between the donor and acceptor atoms.

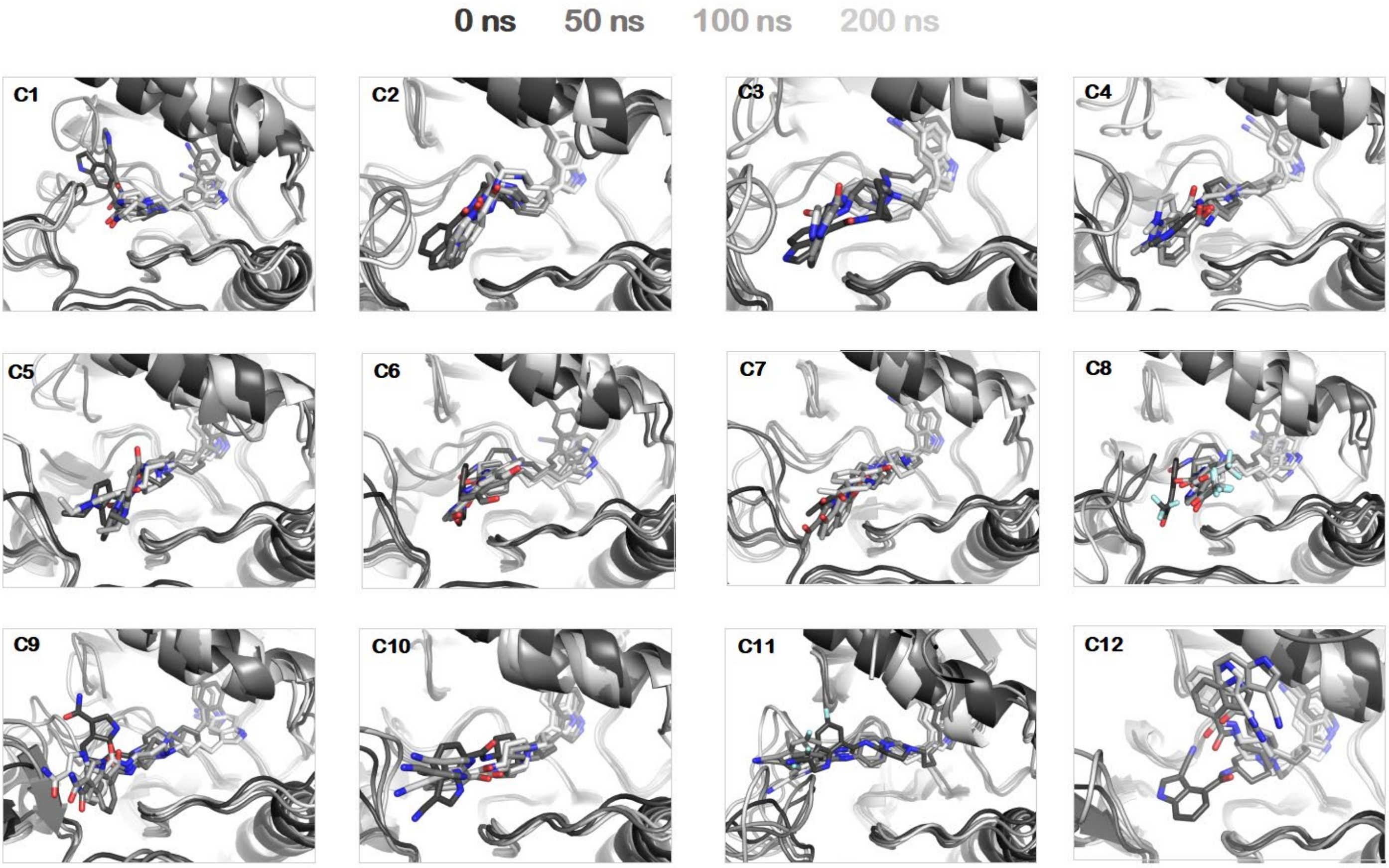


**Figure S2: Structural superimposition of snapshots from molecular dynamics simulation of each complex.** Snapshots correspond to conformations at *t = 0, 50, 100,* and *200* ns, progressing from darker to lighter shades.

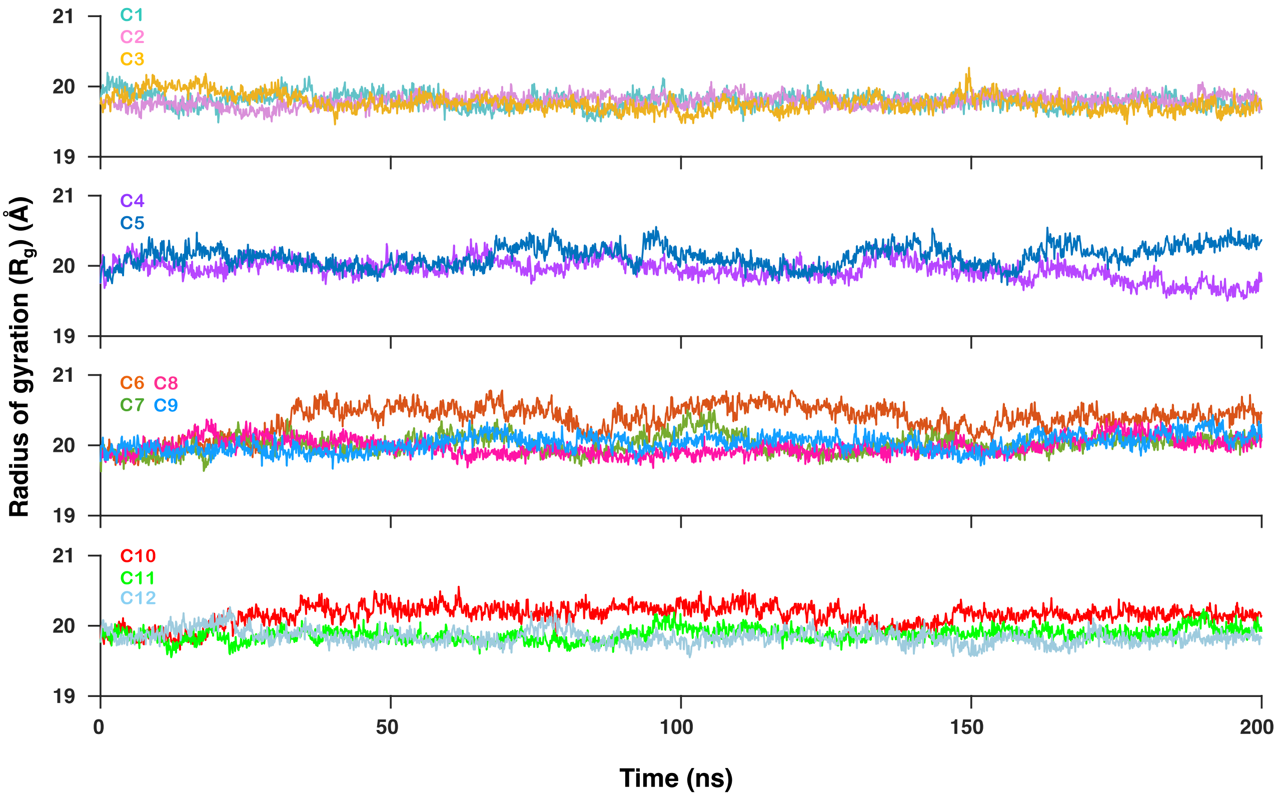


**Figure S3:** The radius of gyration ($R_g$) data for the protein in each protein/inhibitor complex.

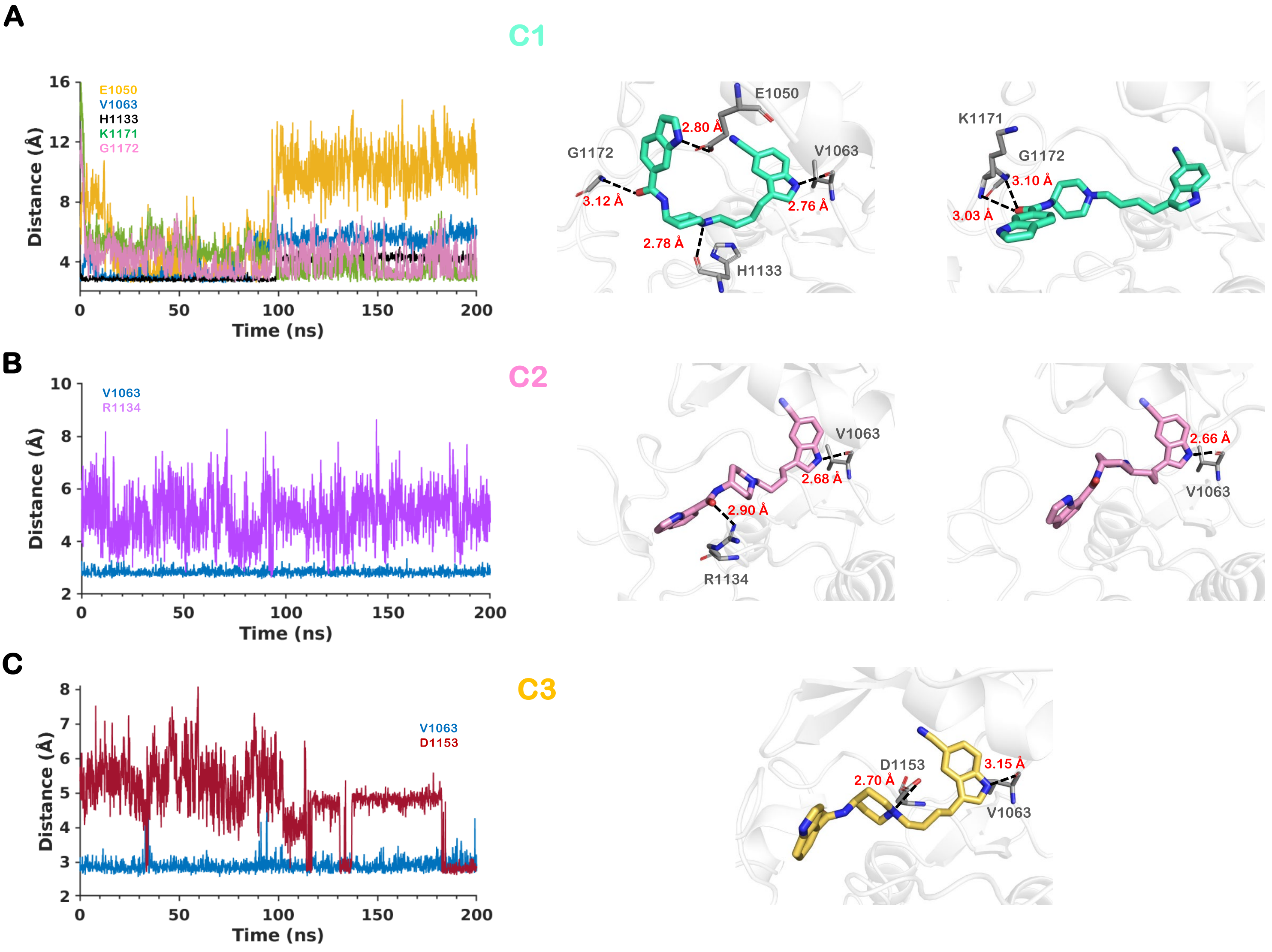


**Figure S4: Time evolution data corresponding to distances for hydrogen bonds formed between the IGF1RK residues and the inhibitor atoms.** Data shown are for Group A inhibitors (C1, C2, and C3). The snapshots are derived from the most populated MD conformation(s) of each complex.

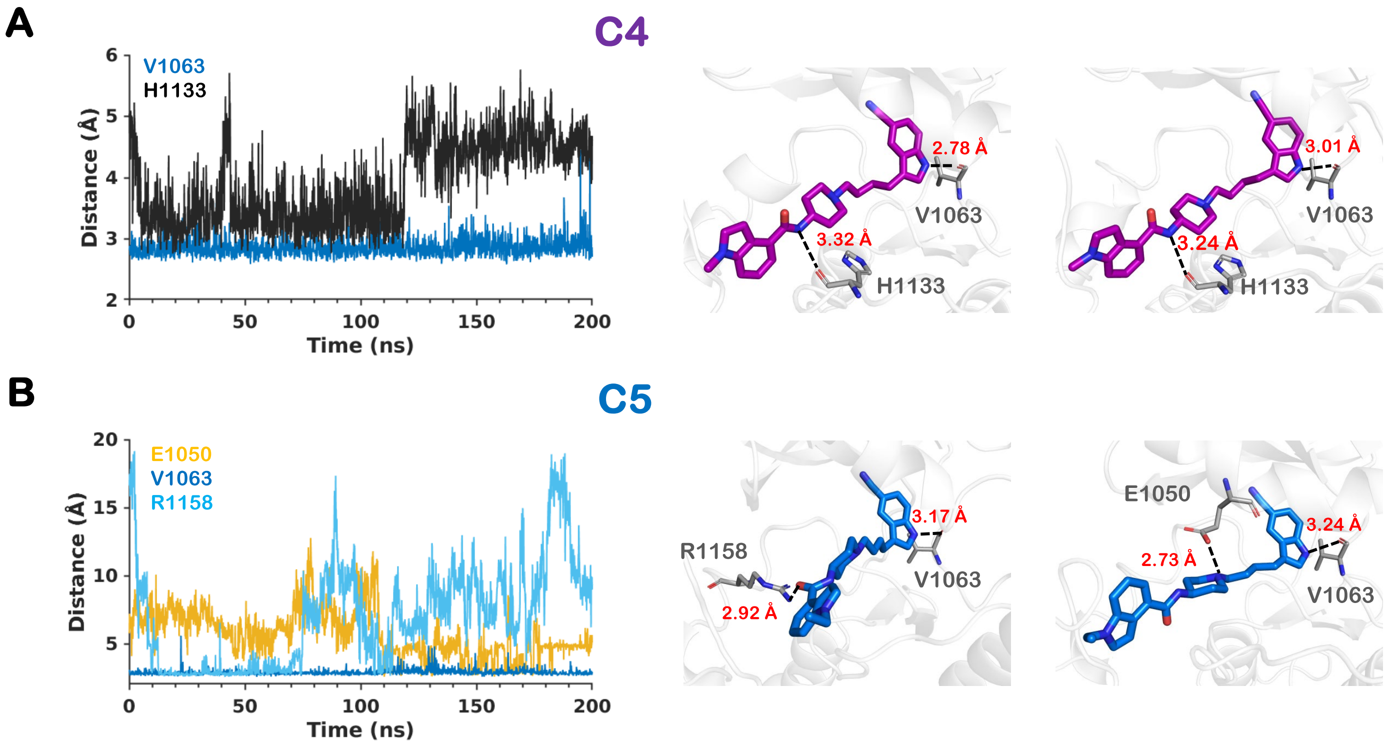


**Figure S5:** Data similar to Figure S4 are shown for Group B inhibitors (C4 and C5).

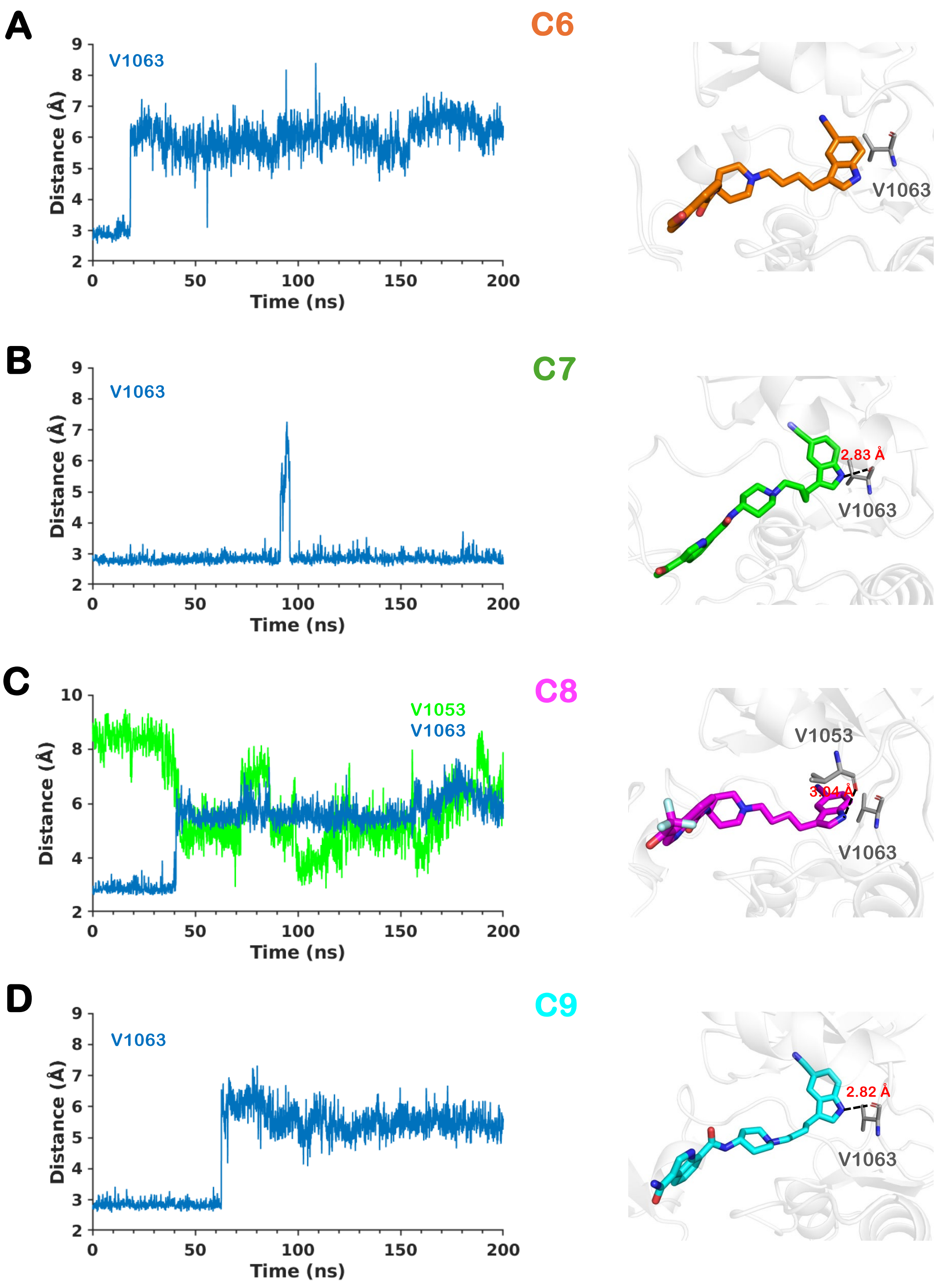


**Figure S6:** Data similar to Figure S4 are shown for Group C inhibitors (C6, C7, C8 and C9).

## C12

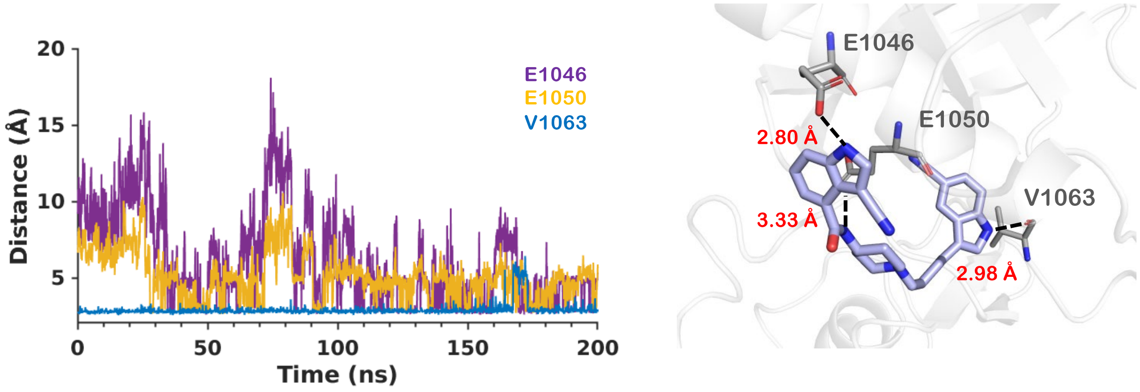


**Figure S7:** Data similar to Figure S4 is shown for a Group D inhibitor (C12).

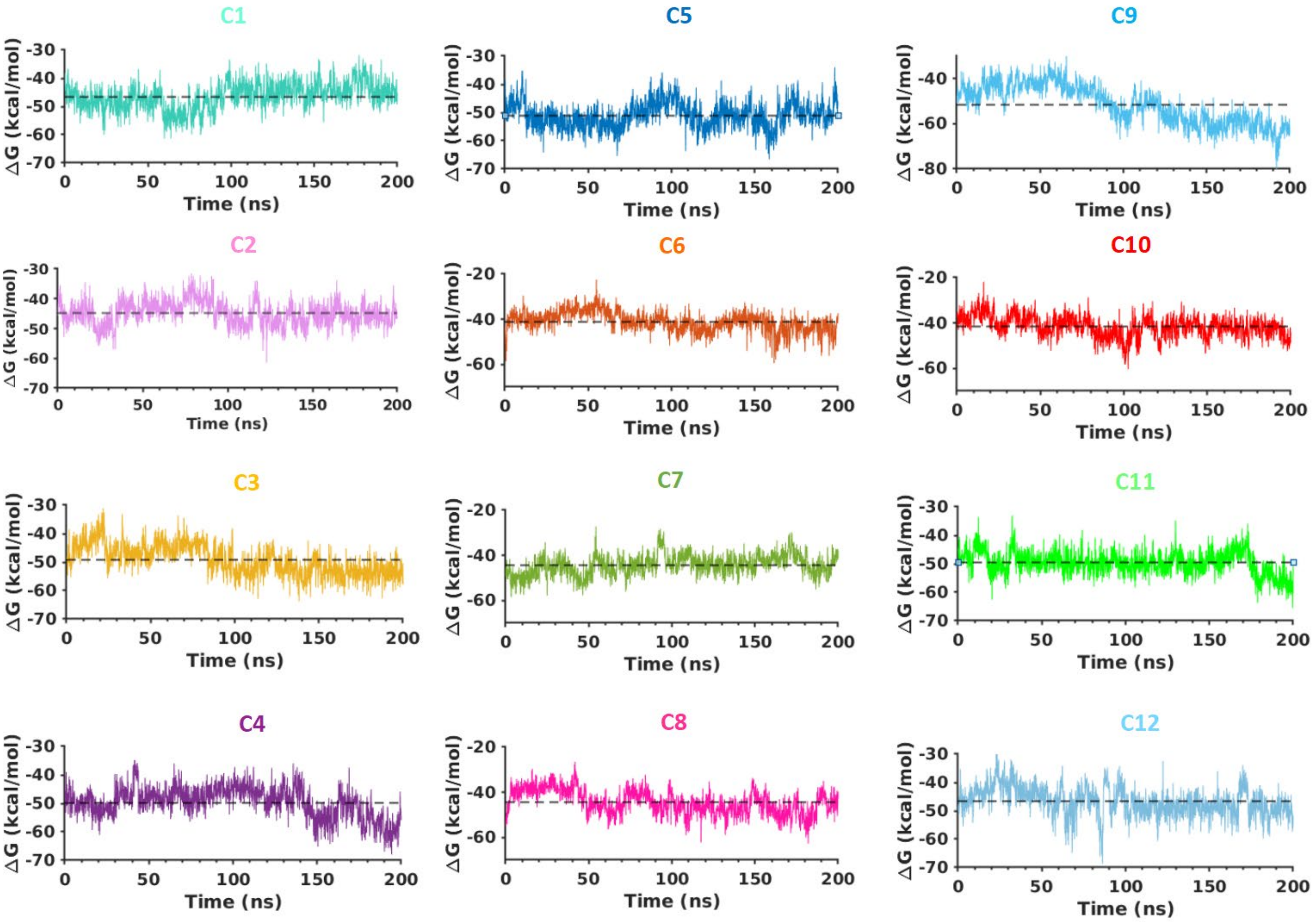


**Figure S8: The time evolution data for the binding free energy (ΔG) of each allosteric inhibitor with the IGF1RK.** The black dashed line in each plot represents the average **Δ**G value.

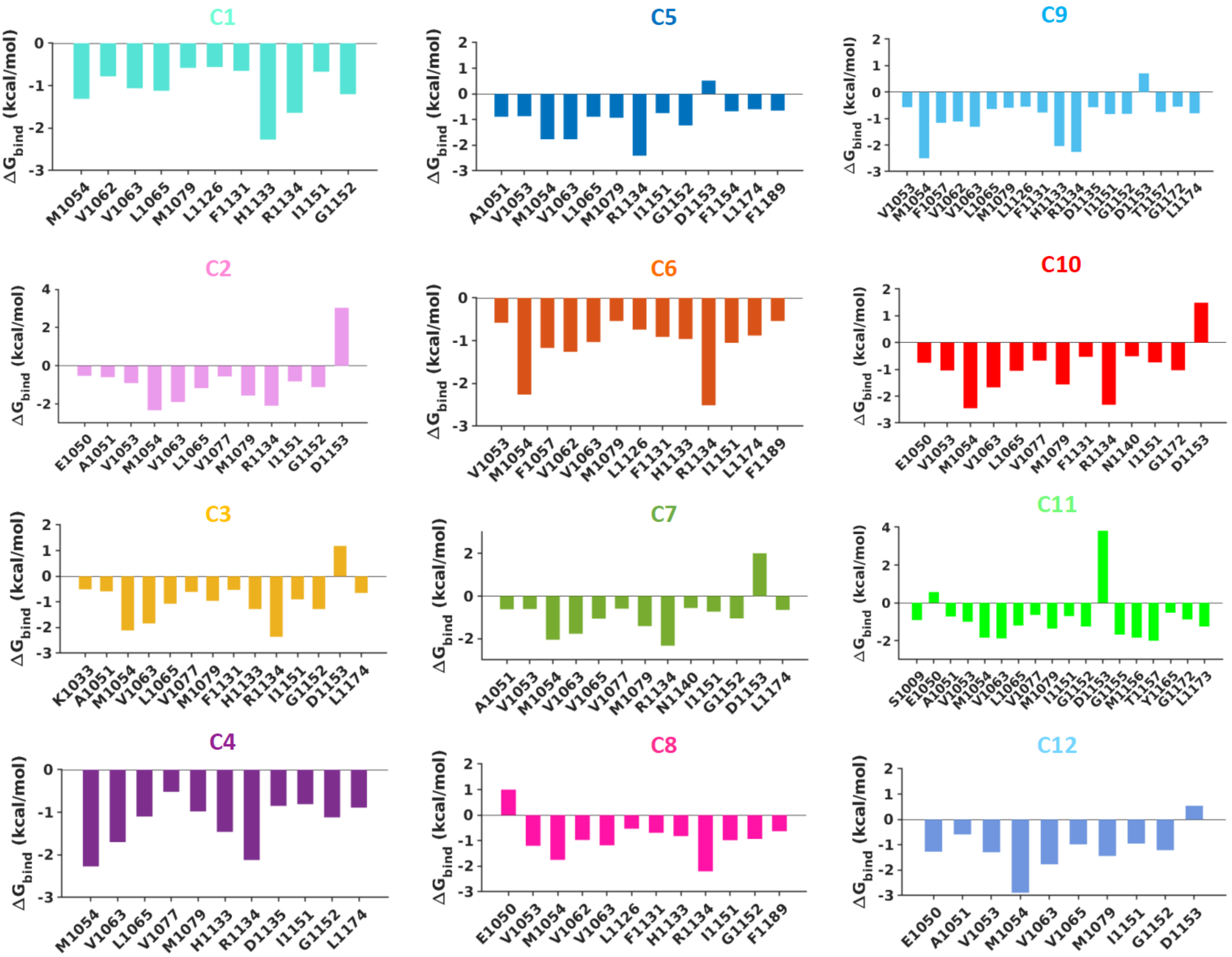


**Figure S9: The residue-level binding free energy contribution for each IGF1RK-inhibitor complex.** The **Δ**$G_{bind}$ for each residue involved in the interaction between the allosteric inhibitor and IGF1RK is shown for each complex.

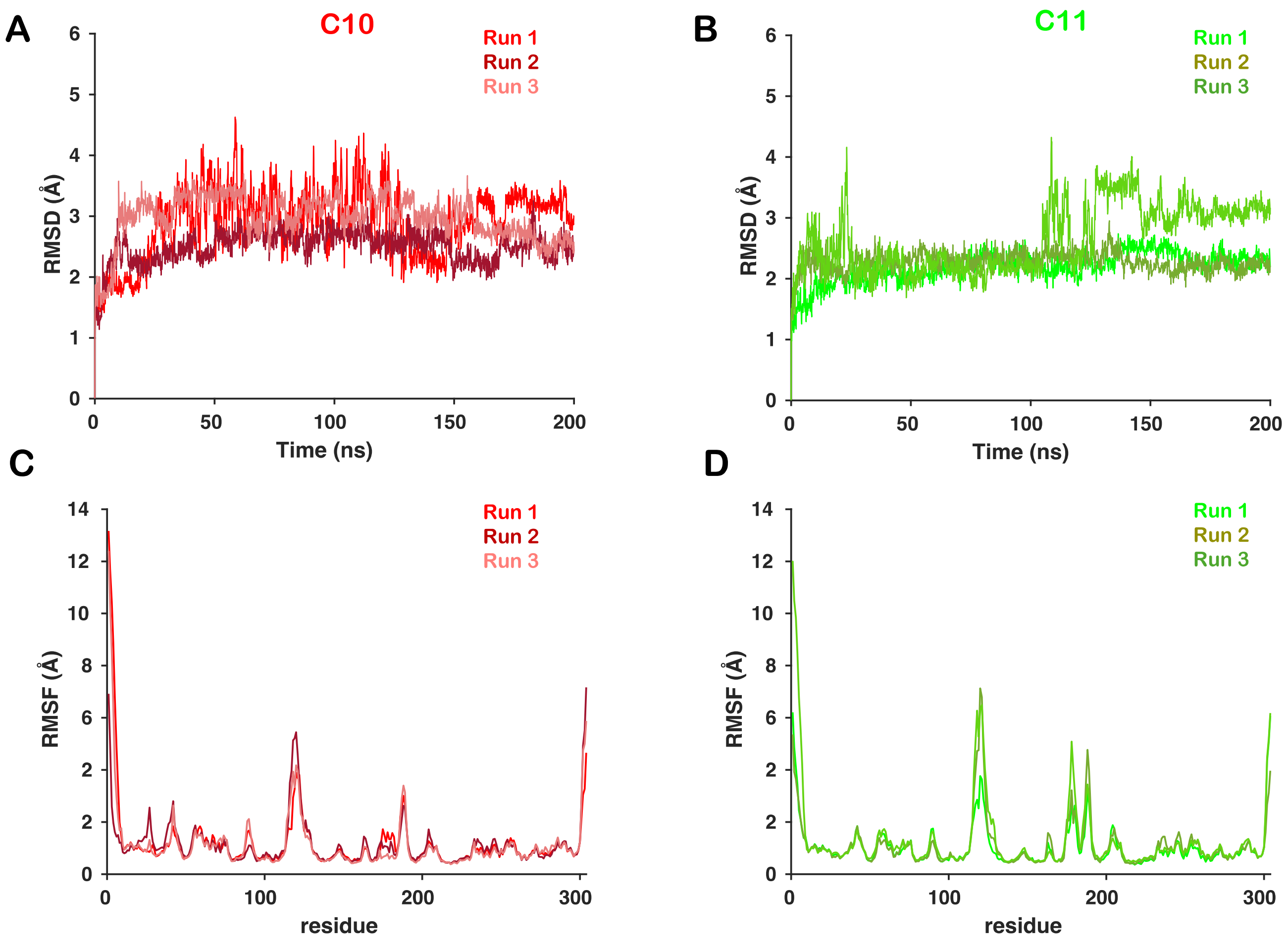


**Figure S10: RMSD and RMSF.** The Root Mean Squared Deviation (RMSD) data for the protein backbone atoms from three independent simulations of complexes of (A) C10 and (B) C11 inhibitors. The Root Mean Squared Fluctuation (RMSF) per residue for the Cα atoms for (C) C10 and (D) C11 inhibitors.
